# Tales of Two α-Carboxysomes: the Structure and Assembly of Cargo Rubisco

**DOI:** 10.1101/2022.03.15.484529

**Authors:** Tao Ni, Yaqi Sun, William Seaton-Burn, Monsour M. J. Al-Hazeem, Yanan Zhu, Xiulian Yu, Lu-Ning Liu, Peijun Zhang

## Abstract

Carboxysomes are a family of bacterial microcompartments in cyanobacteria and chemoautotrophs. It encapsulates carbonic anhydrase and Ribulose 1,5-bisphosphate carboxylase/oxygenase (Rubisco) catalysing carbon fixation inside a proteinaceous shell. How Rubisco packs into the carboxysomes is unknown. Using cryo-electron tomography and subtomogram averaging, we present 3D organization of Rubisco inside two types of native α-carboxysomes from a marine α-cyanobacterium *Cyanobium* sp. PCC 7001 and a chemoautotrophic bacterium *Halothiobacillus neapolitanus*. We determined the structures of Rubiscos within native *Halothiobacillus* and *Cyanobium* carboxysomes at 3.3 Å and 3.8 Å resolution respectively and further identified an associated CsoS2 segment. Interestingly, CsoS2 is only associated with a sub-population of Rubiscos that are close to the shell in *Halothiobacillus*, but with all Rubiscos throughout *Cyanobium* carboxysome. Moreover, Rubiscos in *Cyanobium* carboxysomes are organized in three concentric layers whereas Rubiscos in *Halothiobacillus* carboxysomes are arranged in spiral arrays. Calcium treatment induced a drastic re-organization of Rubiscos, converting these two distinct assemblies into ordered lattice arrays in both α-carboxysomes. Our findings provide critical knowledge of the assembly principles of α-carboxysomes, which may aid in rational design and repurpose of carboxysome structures for new functions.

## Introduction

Bacterial cells have evolved defined internal structures, including intracellular membranes, vesicles, and membrane-less organelles, to compartmentalize and tune metabolic reactions in space and time^1,2^. Bacterial microcompartments (BMCs) are a paradigm of metabolic organelles that are composed purely of proteins and are widespread across the bacterial kingdom^3,4^. By sequestering key enzymes and pathways from the bacterial cytoplasm to enhance catalytic performance and reduce toxicity or unwanted side reactions, BMCs play vital roles in autotrophic CO_2_ fixation and catabolic processes^5,6^.

The first structurally discovered BMCs were carboxysomes, which serve as the central CO_2_-fixing organelles in all identified cyanobacteria and many chemoautotrophs^7-9^. The carboxysome encapsulates carbonic anhydrase and the primary CO_2_-fixing enzyme, ribulose-1,5-bisphosphate carboxylase oxygenase (Rubisco), within a protein shell that structurally resembles a virus capsid^10^. The shell is semi-permeable, ensuring influx of bicarbonate and accumulation of CO_2_ around encapsulated Rubisco to enhance carbon fixation^11,12^. The current models of carboxysome shells are predominantly based on an icosahedral architecture, given the assumption that hexameric proteins form shell facets while pentameric proteins occupy the vertices of the icosahedron^13,14^. However, increasing experimental evidence has highlighted the structural variability and plasticity of BMC shells^15-19^.

Rubisco is among the most abundant components of carboxysomes^15,17^. How Rubisco enzymes are organized within the carboxysome to conduct efficient carboxylation has been a long-standing question. Carboxysomes can be divided into two lineages, α- and β-carboxysomes, which differ in the forms of Rubisco and their structural protein composition. It was shown that the internal organization of β-carboxysomes from freshwater β-cyanobacteria is highly packed with paracrystalline arrays of Rubisco^20^. This packaging, mediated by the scaffolding protein CcmM, results in formation of a liquid-like condensate^21^, which subsequently triggers shell encapsulation and eventually construction of full β-carboxysomes^22,23^. In contrast, the Rubisco packing and biogenesis of α-carboxysomes remain unclear.

The α-carboxysome components are encoded by genes mainly in a *cso* operon in the genome. The shell is constructed by CsoS1 hexameric proteins and CsoS4 pentamers. The intrinsically disordered protein CsoS2 serves as the linker bridging the shell and the cargo Rubisco. The N-terminus of CsoS2 binds with Rubisco and induces Rubisco condensation^24^; while the C-terminus of CsoS2 is presumed to interact with shell proteins^25,26^. Despite a few cryo-electron tomography (cryoET) studies on α-carboxysome structures^27-30^, the details of Rubisco structure and its assembly within the intact α-carboxysome and biogenesis of α-carboxysomes remain unclear.

Here we investigated the structure and assembly of Rubiscos in two representative α-carboxysomes from a marine α-cyanobacterium *Cyanobium* sp. PCC 7001 and a chemoautotrophic bacterium *Halothiobacillus neapolitanus* (hereafter *Cyanobium* and *Halo*, respectively). Using cryoET and subtomogram averaging (STA)^31^, we determined the structures of Rubisco within these native α-carboxysomes at neat-atomic resolution and identified the associated domain of CsoS2. Interestingly, whereas Rubisco and CsoS2 association was observed throughout the carboxysome from *Cyanobium*, CsoS2 was found to only associated with the outer-shell Rubiscos in the *Halo* α-carboxysome. Furthermore, while Rubiscos are organized in concentric shells in *Cyanobium* α-carboxysomes, they form intertwining spirals in *Halo* carboxysomes, and intriguingly, both rearrange into a higher-order assembly upon Ca^2+^ treatment. The results advance our knowledge about Rubisco organization and protein interactions within the α-carboxysomes, which may aid in rational design and repurpose of carboxysome structures for new functions.

## Results

### Structure and assembly of Rubisco in Cyanobium carboxysomes

CryoEM images show *Cyanobium* α-carboxysomes is relatively homogeneous in size (Fig. 1a), which prompted us to attempt its structural determination using single particle cryoEM (SPA). However, 2D class averages suggest structural variation of carboxysomes (Fig. 1b). Further 3D classification of *Cyanobium* α-carboxysomes only yielded a low-resolution map from a subset of 2D classes with C1 symmetry (32%), which shows polyhedron with 20 faces and 12 vertices but deviate from an icosahedron (Fig. 1c). Interestingly, Rubisco densities are arranged in three concentric layers which are separated by 11 nm (Fig. 1d). The individual Rubiscos, however, were not resolved. This variable morphology of *Cyanobium* carboxysomes is confirmed by cryoET (Fig. 1e).

**Figure 1.**
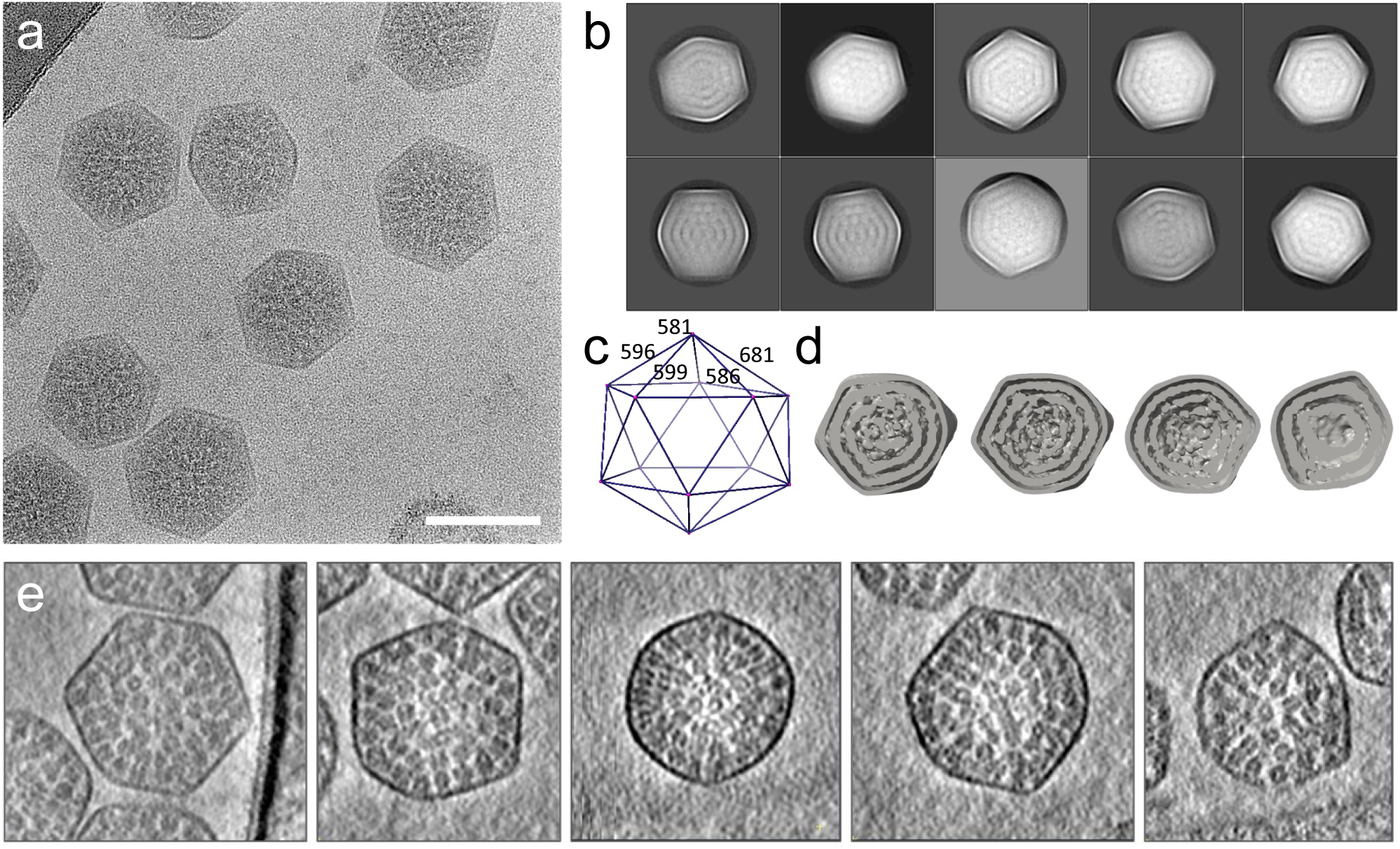
CryoEM SPA of *Cyanobium* carboxysomes. (a) A representative micrograph of *Cyanobium* carboxysomes. (b) 2D class averages of *Cyanobium* carboxysomes. (c-d) Reconstruction of *Cyanobium* carboxysomes without symmetry, shown in geometry (c) and cross-sections (d). (e) A gallery of non-icosahedral *Cyanobium* carboxysomes (central tomographic slices) with variable size and shape. Scale bar 100 nm.

To determine the structure of Rubisco within the intact carboxysomes and analyse its organization, we performed cryoET STA using emClarity^32^. The individual Rubisco can be readily delineated in the raw tomograms (Fig. 2a, Movie 1). Template matching and mapping back the position and orientation of individual Rubisco compelxes in the original tomograms revealed three concentric layers of Rubiscos that are oriented with their 4-fold axis along the radial direction (Fig. 1b-c). The radial distances of three shells are peaked at 208Å, 308 Å and 413 Å (Fig. 2c left) and angles of ∼15° from radial axis (Fig. 2c right). Interestingly, this concentric shell arrangement of Rubiscos was disrupted upon Ca^2+^ treatment (Fig. 1d-f). Instead, Rubiscos are rearranged into extended 3D arrays (Fig. 2d-e, Movie 2). While treatment with K^+^ and Mg^2+^ did not yield such an effect, the mechanism of this remarkable Ca^2+^-induced reorganization of Rubisco is still not clear and requires further investigation.

**Figure 2.**
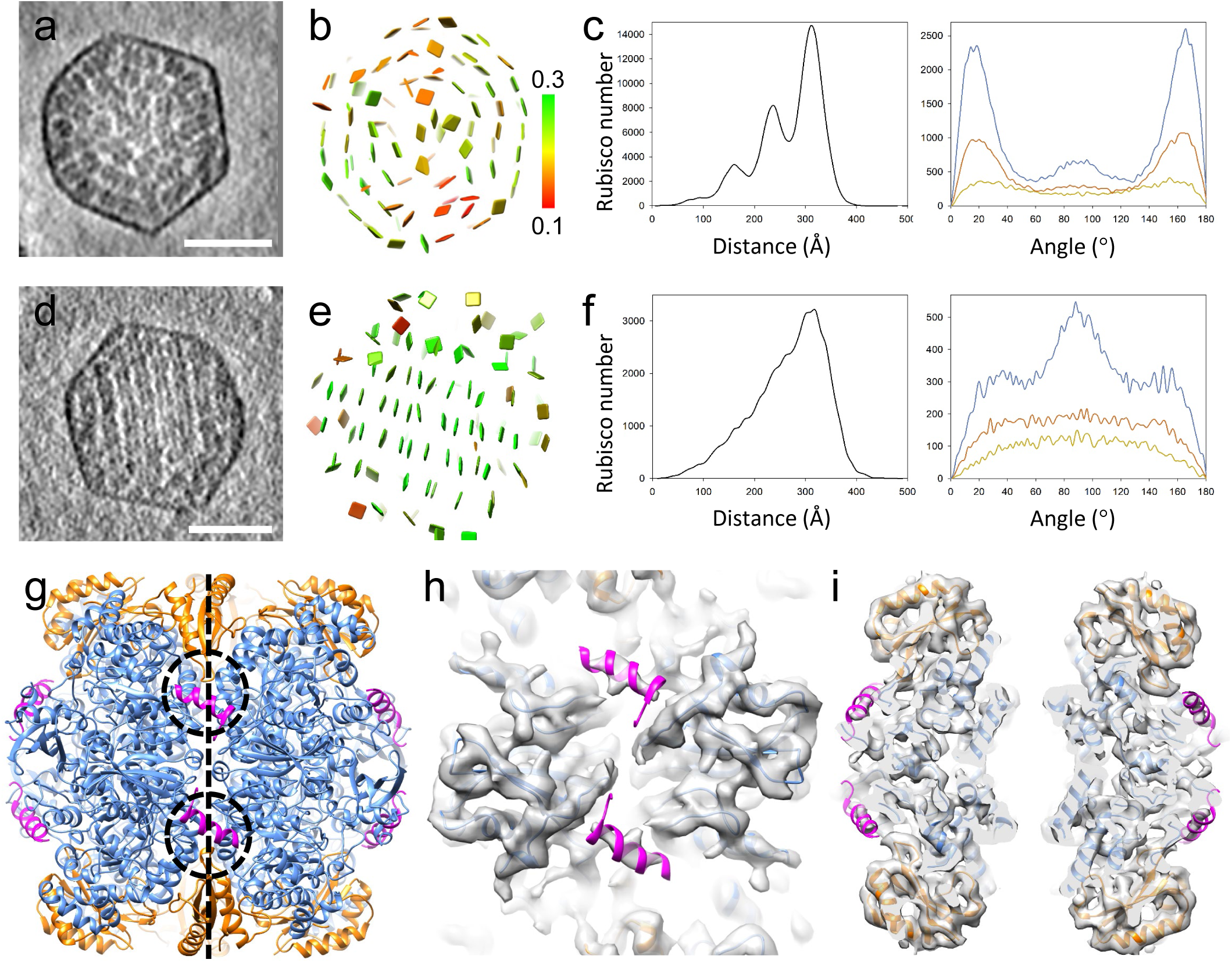
Structure and organization of Rubisco within intact *Cyanobium* carboxysomes. (a) A tomograms slice (26.8 A thickness) of a *Cyanobium* carboxysome. (b) The position and orientation of individual Rubisco is mapped back to the tomogram of carboxysome, shown as a discs normal the 4-fold symmetry axis of Rubisco and colored according to the cross-correlation values between individual Rubisco and the STA map. (c) Radial (left) and angular (right) distributions of Rubisco in *Cyanobium* carboxysomes. (d) A tomograms slice (26.8 A thickness) of a Ca^2+^ treated *Cyanobium* carboxysomes. (e) The individual Rubisco is mapped back to the tomogram of a Ca^2+^ treated carboxysome. (f) Radial (left) and angular (right) distributions of Rubisco in Ca^2+^ treated carboxysomes. (g-i) CryoET STA structure of the Rubisco in *Cyanobium* carboxysomes at 3.8 Å resolution, shown in atomic model (g), a top slice (h) and a central slice (i) overlapped with density. CbbL and CbbS are in blue and gold, respectively. The CsoS2 peptide density was resolved and modelled in magenta. Dashed circles indicate the interaction between CsoS2 and CbbS. Dashed line indicates the 4-fold axis. Scale bars, 100 nm.

Further cryoET STA of Rubiscos resulted a density map at 3.8 Å resolution, unprecedent for the *in situ* Rubisco structure (Fig. 2g, Fig. S1a, Movie 3). Since there is no atomic model for the *Cyanobium* Rubisco, we built an MDFF model based on alphafold2 prediction^33^ (Fig. 2g). The overall structure of *Cyanobium* Rubisco hexadecamer is very similar to those published homologues, with an RMSD of 0.86 Å between this and the *Halo* Rubisco crystal structure (1SVD). Surprisingly, we observed an additional density that is not part of Rubisco (Fig. 2h-i). This density matches very well to the helical peptide of CsoS2 (Fig. 2h-i, magenta), as in the crystal structure of *Halo* Rubisco in complex with the peptide (PDB: 6UEW)^34^. CsoS2 serves as a linker connecting the carboxysome shell using its C-terminal region to Rubisco through its N-terminus^34,35^. To determine whether the CsoS2 interacts with the Rubiscos in all three concentric layers, we obtained STA maps Rubisco from three shells separately. All three maps display the density corresponding to the CsoS2 N-terminal peptide, indicating its essential role in packaging Rubisco in the *Cyanobium* α-carboxysome (Fig. S2).

### Structure and assembly of Rubisco in Halo carboxysomes

To understand how Rubiscos are organized in different α-carboxysomes and whether there is a conserved architecture, we analyzed a distant α-carboxysomes, the *Halo* α-carboxysome^36^. Visual inspection of the tomographic reconstructions revealed that the organization of Rubiscos within *Halo* carboxysomes differs from those within *Cyanobium* carboxysomes: *Halo* Rubiscos form intertwined spirals instead of concentric layers (Fig. 3a, red arrow, Movie 4-5). Compared to the average number of 224 ± 26 Rubiscos contained in *Cyanobium* carboxysomes, there are 274 ± 72 Rubiscos in *Halo* carboxysomes (Fig. S3a-b), which is slightly smaller than the stoichiometric composition determined by the QconCAT-based quantitative mass spectrometry^15^. The distances between two neighbour Rubiscos in both carboxysomes are very similar (Fig. S3c-d).

**Figure 3.**
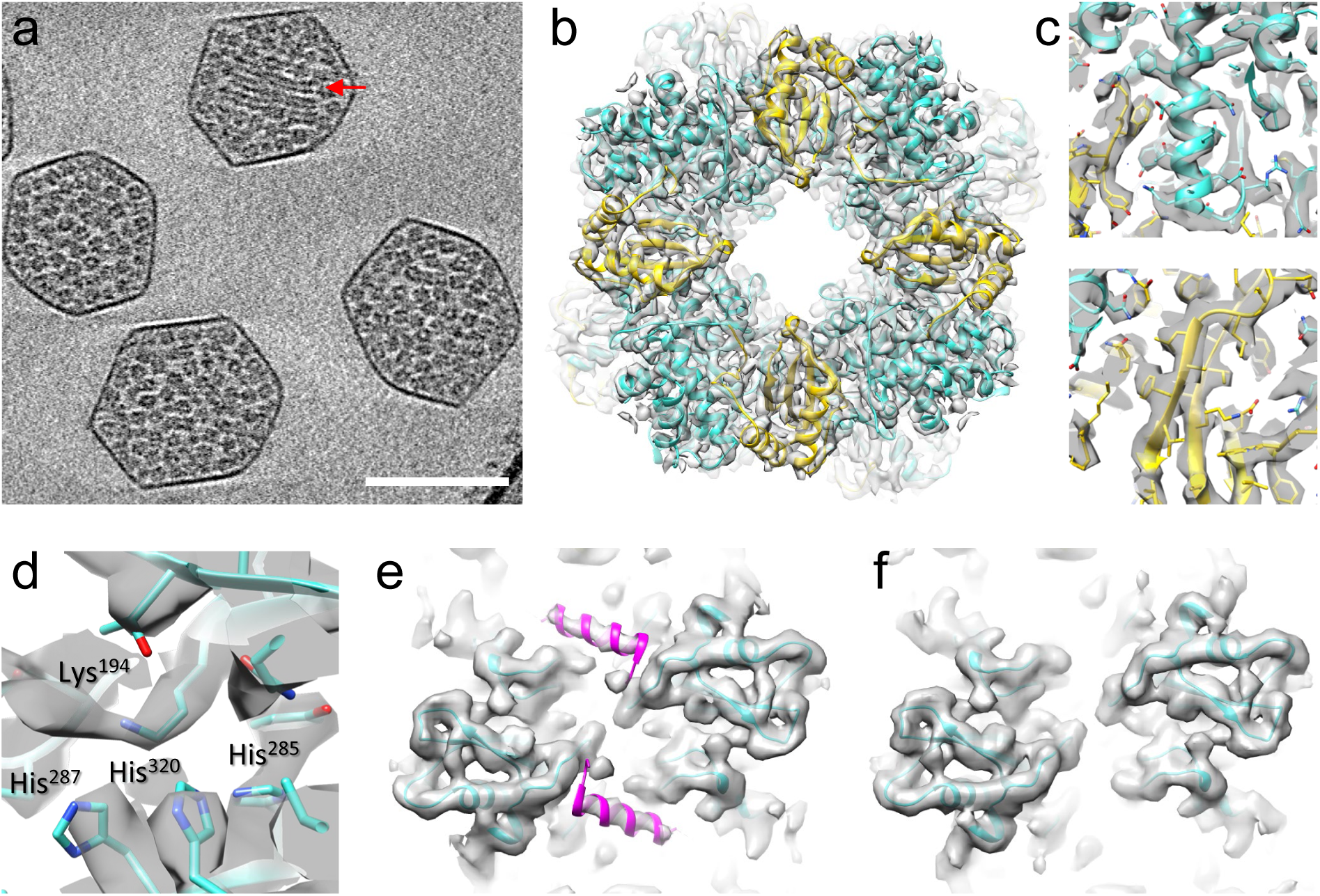
CryoET STA structure of Rubisco within intact *Halo* carboxysomes. A tomograms slice (33.5 A thickness) of a *Halo* α-carboxysome. Strings of Rubisco are marked by the red arrow. (b) CryoET STA structure of Rubisco at 3.3 Å, overlapped with the real-space refinned atomic model of its components, CbbL (cyan) and CbbS (yellow). (c) Details of Rubisco density map and the atomic model shown with side-chains. (d) The Rubisco catalytic site. (e) CryoET STA structure of Rubiscos close to outer shell, overlapped with the atomic model. The CsoS2 peptide density was resolved and modelled in magenta. (f) CryoET STA structure of Rubiscos within 300 Å from the center, overlapped with the atomic model. Scale bar, 100 nm.

CryoET STA of Rubiscos in *Halo* carboxysomes resulted in a density map at 3.3 Å resolution, which allows a real-space refinement of the *in situ* Rubisco structure (Fig. 3b-c, Fig. S1b, Movie 6). There is little deviation between the refined cryoET STA structure and the crystal structure of *Halo* Rubisco (PDB: 1SVD) (RMSD of 0.35 Å). The carbamylation of Lysine 194 in the catalytic site is clearly resolved, together with three key histidine residues, likely important for regulating Rubisco activity^37^ (Fig. 3d). Intriguingly, unlike the extra density identified in the *Cyanobium* Rubisco map, we observed no additional density corresponding to the CsoS2 peptide. We reasoned that this might be due to a lower overall occupancy of CsoS2 with *Halo* Rubisco and speculate that CsoS2 might have distinct associations with certain Rubisco populations. We, therefore, obtained STA maps of Rubisco from those close to shell and those within 30 nm from the center, separately. Remarkably, there is a clear density corresponding to the CsoS2 helical peptide in the Rubiscos adjacent to the shell, but is absent in the Rubiscos near the center (Fig. 3e-f).

In 38% of *Halo* carboxysomes, Rubiscos are organized in a spiral array (Fig. 4a, Fig. S4a, Movie 4-5), which accounts for ∼8% of total Rubiscos in these carboxysomes. The Rubisco spiral array tends to localize in the centre of carboxysome. The number of Rubisco strings varies among individual carboxysomes from 2 to 35 (mean ± SD = 12 ± 6, Fig. S4b), and the lengths of them also vary from 2 to 9 Rubiscos (mean ± SD = 5 ± 2, Fig. S4c). The spiral array is formed by near-parallel packing of Rubisco strings: the central Rubisco string is surrounded by 6 strings (Fig. 4b, Movie 7, Fig. S3a). To understand the molecular interactions between the Rubiscos in the string-like assembly, we further determined the Rubisco dimer structure at 4.2 Å resolution using cryoET STA and docked the atomic model of *Halo* Rubisco (Fig. 4c, Fig. S1c). As shown in Figure 4c-d, the Rubisco tandem dimer interface is primarily mediated by four CbbS subunits, providing charge-charge interactions similar to those observed in the crystal packing (PDB: 1SVD) (Fig. 4d). But the Rubisco tandem dimer in the string assembly is rotated about 7.3° with respect to each other. As with *Cyanobium* carboxysomes, we tested Ca^2+^ effect on the assembly of *Halo* Rubisco. Ca^2+^ treatment induced remarkable re-organization of Rubisco into an extended well-ordered 3D lattice array inside *Halo* carboxysomes (Figure 4e, Movie 8). The propensity of rearrangement is Ca^2+^ concentration dependent and reversible upon Ca^2+^ removal (Fig. S5), whereas other ions, Mg^2+^, Na^+^, K^+^, have no effect (Fig. S6). Mapping back the refined positions and orientations of Rubiscos reveals that the Rubiscos are packed against each other with two interaction interfaces: one along the 4-fold axis and the other normal to the 4-fold axis (Fig. 4f-g). To gain further insights of Ca^2+^-induced Rubisco ordered array assembly, we performed cryoET STA of the Rubisco array subunits containing Rubisco tandem dimers along or normal to the 4-fold axis in the Ca^2+^ treated *Halo* carboxysomes. The STA maps of tandem dimers reveal the former dimer interface similar to the interface identified in the native state (Fig. 4h) and the latter dimer interface likely mediated by the CbbL (Fig. 4i).

**Figure 4.**
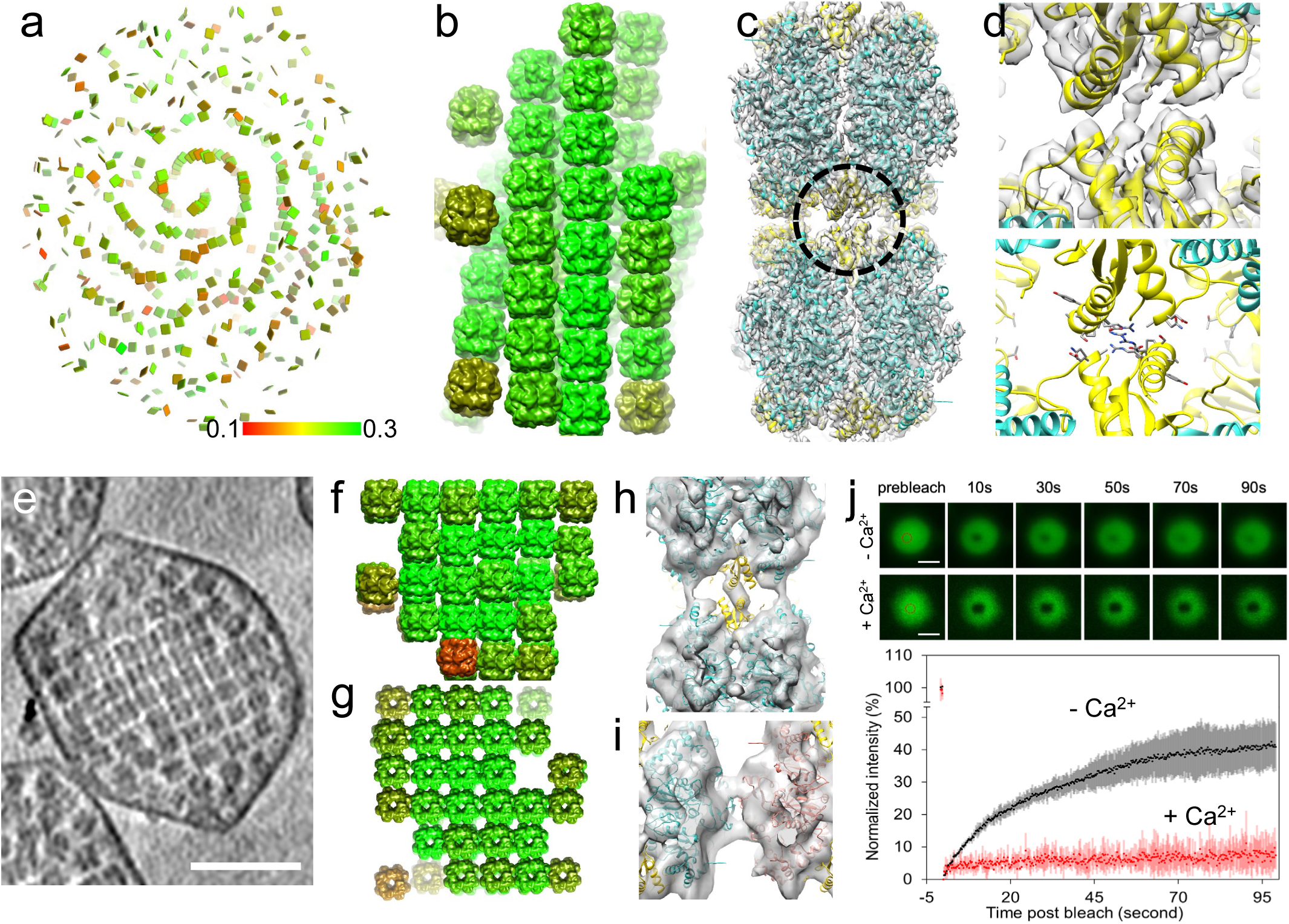
Organization of Rubisco within *Halo* α-carboxysomes. (a) The position and orientation of individual Rubisco is mapped back to the tomogram of a *Halo* carboxysome. Rubisco subvolumes are mapped back to a *Halo* carboxysome. (c) CryoET STA structure of Rubisco dimer stacking along the 4-fold symmetry axis, overlapped with fitted atomic model. (d) A detailed view of the dimer interface between CbbS subunits (circled in c) (top) and shown with charged interface residues (bottom). (e) A tomographic slice of a Ca^2+^ treated *Halo* carboxysome containing a Rubisco lattice array. Scale bar, 50 nm. (f-g) Rubisco subvolumes are mapped back to a Ca^2+^ treated *Halo* carboxysome, viewed normal to (f) and along (g) 4-fold axis. (h-i) CryoET STA of Rubisco dimers along the 4-fold axis (h) and normal to the 4-fold axis (i). (j) FRAP of Rubisco condensates formed by unlabelled *Halo* Rubisco (1.6 μM) and CsoS2-NTD-sfGFP (2 μM). Representative condensates with and without Ca^2+^ treatment (250 mM) are shown before and after bleaching. The sites of bleaching are marked by dashed circles. Scale bar, 2 μm. The change in fluorescence was analysed as a function of time and the *t*_1/2_ of fluorescence recovery is indicated as mean ± s.d. (*n* = 3).

The formation of such highly ordered Rubisco arrays upon Ca^2+^ treatment raises a question regarding Rubisco dynamics inside a carboxysome. To test this, we analysed the assembly dynamics of Rubisco-CsoS2-NTD using fluorescence microscopy. Rubisco–CcmM and Rubisco–CsoS2 form liquid-like matrices, important for carboxysome assembly^21,34^. Combining Rubisco and CsoS2-NTD fused with super-fold GFP (sfGFP) induced formation of round fluorescent condensates (Fig. 4j), characteristic of liquid droplets. Formation of Rubisco condensates in the presence of Ca^2+^ (250 mM), the same condition that triggers highly ordered Rubisco packing within α-carboxysomes, appeared less efficient than without Ca^2+^, as reflected by weaker fluorescence condensates observed. Fluorescence recovery after photobleaching (FRAP) of the droplets showed a notably slower rate of recovery with Ca^2+^ treatment (*t*_1/2_ ≈ 50 s) than without Ca^2+^ treatment (*t*_1/2_ ≈ 20 s, Fig. 4c), demonstrating that Rubiscos in the Ca^2+^-induced ordered packing are more stable. The dynamic nature of the Rubisco–CsoS2-NTD assemblies suggested that weak interactions between Rubiscos, which are salt sensitive, may play roles in liquid-liquid phase separation of Rubisco assemblies.

## Discussion

Understanding the assembly mechanism of carboxysomes is key for its biotechnological applications using synthetic biology. In this work we present the 3D organization of Rubiscos inside two native α-carboxysomes from a marine α-cyanobacterium *Cyanobium* sp. PCC 7001 and a chemoautotrophic bacterium *Halothiobacillus neapolitanus*. Subtomogram averaging of Rubisco compelxes at 3.3 and 3.8 Å resolution reveals Rubisco in resting state inside native carboxysomes. This allows further investigation of Rubisco assembly and functional regulation within the intact carboxysome. It also provides approaches to study other bacterial microcompartments like β-carboxysomes and metabolosomes. We uncovered different organization of Rubiscos in two α-carboxysomes: Rubisco in the *Cyanobium* carboxysome are organized in concentric layers along the shell, with the Rubisco small subunit CbbS facing towards the shell; Rubiscos in the *Halo* carboxysomes are arranged in spiral arrays, with the principal interaction mediated by 4 pairs of CbbS between the neighbours. Several reasons could contribute to the difference in Rubisco organization within these two α-carboxysomes in their native states. Firstly, the Rubiscos surface display different surface electrostatic property (Fig. S7): *Halo* Rubisco display charged surface in the CbbS, which promote the assembly of string-like structures in the spiral array; in contrast, CbbS of *Cyanobium* Rubisco shows largely positively-charged surface, possibly repelling the inter-molecular interaction along the 4-fold axis (Fig. S7). Secondly, CsoS2 in these two carboxysomes are more divergent than Rubisco (37% identity in CsoS2, 81.7% in CbbL and 50% in CbbS). *Halo* carboxysomes possess two isoforms of CsoS2, translated via programmed ribosomal frame shifting^38^, with one as truncated form lacking the C-terminal region responsible for carboxysomal shell anchoring. However, *Cyanobium* carboxysome only contain the full-length form of CsoS2.

Our results provide the direct evidence for the CsoS2-Rubisco interaction within native α-carboxysomes. The patterns of CsoS2-Rubisco interaction, however, are distinct between two α-carboxysomes. In *Cyanobium* carboxysomes, CsoS2 binds to all Rubiscos across three concentric layers (Fig. 2h-i). In contrast, in *Halo* carboxysomes, CsoS2 primarily associates with Rubiscos that are close to the shell (Fig. 3e-f). The divergence of CsoS2 may contribute, in part, to the differences in CsoS2-Rubisco interaction and subsequently Rubisco packing and dynamics within the carboxysome, which likely correlate with Rubisco activity and biogenesis/repair. The fact that in both α-carboxysomes Rubiscos close to the shell are connected to CsoS2 suggests that Rubiscos are potentially recruited and encapsulated via CsoS2 linkage for the initial assembly, distinct from the packing and biogenesis of β-carboxysomes. Furthermore, Ca^2+^-treatment induces remarkable re-organization of Rubiscos into ordered 3D arrays in both α-carboxysomes, suggesting a potentially conserved mechanism, the details of which merit further investigation.

## Competing interests

The authors declare that they have no competing interests.

## Acknowledgments

We are grateful to Yuriy Chaban and Yuewen Sheng for technical support. We thank Prof. Ian Prior and Mrs. Alison Beckett at the Liverpool Biomedical Electron Microscopy Unit for support of electron microscopy. We acknowledge Diamond for access and support of the cryoEM facilities at the U.K. national eBIC (proposals NT21004 and EM20223), funded by the Wellcome Trust, MRC, and BBSRC. This work was supported by Wellcome Trust Investigator Award 206422/Z/17/Z, the UK Biotechnology and Biological Sciences Research Council grant BB/S003339/1, the ERC AdG grant (101021133), the National Natural Science Foundation of China (32070109), the National Key R&D Program of China (2021YFA0909600), the Royal Society (URF\R\180030, RGF\EA\181061, RGF\EA\180233), the Biotechnology and Biological Sciences Research Council Grant (BB/V009729/1, BB/M024202/1), the Leverhulme Trust (RPG-2021-286). The research was supported by the Wellcome Trust Core Award Grant Number 203141/Z/16/Z with additional support from the NIHR Oxford BRC.

## Contributions

T.N., L-N.L. and P.Z. conceived the projects; Y.S. and M.M.J.A performed carboxysome preparation and biochemical characterization; T.N. made cryoEM samples; T.N. and Y.Z. collected cryoEM SPA and cryoET datasets; T.N. and W.S. analyzed cryoEM SPA data; T.N. processed cryoET data and performed subtomogram averaging with assistance of W.S, Y.Z. and X.Y.; T.N. and Y.Z refined the Rubisco structures. T.N and P.Z. analyzed the Rubisco structures and organization; Y.S. performed photo-bleaching and Ca^2+^ treatment experiments; T.N., L-N.L. and P.Z. wrote the manuscript with support from all authors.

## Data availability

All data needed to evaluate the conclusions in the paper are present in the paper and/or the Supplementary Materials. The cryoET subtomogram averaging density maps and corresponding atomic models have been deposited in the EMDB and PDB, respectively. The accession codes are listed as follows: *Cyanobium* Rubisco from all the carboxysome Rubisco (PDB: XXX and EMD-XXX), *Cyanobium* Rubisco from outer layer (EMD-XXX), inner layers (EMD-XXX); *Halothiobacillus* Rubisco inside carboxysomes (PDB: XXX and EMD-XXX), close to shell (EMD-XXX), within 300 Å from the carboxysome center (EMD-XXX).

## Methods

### Purification of H. neapolitanus and carboxysomes and calcium treatments

The *Halothiobacillus neapolitanus* (*H. neapolitanus*) strain used in this work was acquired from ATCC (The American Type Culture Collection). Cell cultivation and carboxysome purification were performed as described previously ^39^. Seeding cells were maintained in liquid ATCC medium 290 or on ATCC 290 1.5% agar plates and inoculate in the Vishniac and Santer medium ^40^ in a 5-liter fermenter (BioFlo 115, New Brunswick Scientific, US) and were kept at constant pH 7.6 through supplement of 3 M KOH. The growth was maintained at 30°C with agitation kept at 250-300 rpm. The air supply that set at 500 L.min^-1^ for initial growth and reduced to 200 L min^-1^ 24-48 hours prior to cell collection. Cells were pelleted by sequential centrifugation at 12,000 g for 10 min, 300 g for 15 min and 12,000 g for 10 min in TEMB buffer (10 mM Tris-HCl, pH 8.0, 10 mM MgCl_2_, 20 mM NaHCO_3_, 1 mM EDTA). Cells were treated by egg lysosome (at a final concentration of 0.5 mg mL^-1^) for 1 hour at 30°C, and then disrupted via glass beads beating (150-212 μm glass bead, acid washed, Sigma-Aldrich, US). The lysates were further treated with 33% (v/v) B-PERII (ThermoFisher Scientific, UK) and 0.5% (v/v) IGEPAL CA630 (Sigma-Aldrich, US). Crude carboxysome enrichment was pelleted at 48,000 g, resuspended, and then loaded to a step sucrose gradient (10-60%) for a 35-min centrifugation at 105,000 g. The milky layer of enriched carboxysome was harvested, and sucrose was removed by an additional round of ultracentrifugation after dilution with TEMB buffer. The final pure carboxysome pellet was resuspended in a small volume of TEMB buffer. Unless indicated otherwise, all procedures were performed at 4°C.

*Cyanobium* sp. PCC 7001 (Pasteur Culture Collection of Cyanobacteria, PCC) cells were grown in 4 L of BG-11 medium under constant illumination at 30°C with constant stirring and bubbling with air. Carboxysomes were purified as described previously ^41^. Cells were collected by centrifugation (6000 g, 10 min) and resuspended in TEB buffer (5 mM Tris-HCL, pH 8.0, 1 mM EDTA, 20 mM NaHCO_3_) with additional 0.55 M mannitol and 60 kU rLysozyme (Sigma-Aldrich, United States). Cells were then incubated overnight (20 h) with gentle shaking at 30°C in the dark, and were collected via centrifugation (6000 g, 10 min). Cells were placed on ice and resuspended in 20 mL ice-cold TEB containing an additional 5 mL 1 µm Silicone disruption beads. Cells were broken via bead beating for 8 min in one-minute intervals of vortex, and 1 min on ice. Broken cells were separated from the beads, and the total resuspension volume was increased to 40 mL with TEB buffer containing an additional 4% IGEPAL CA-630 (Sigma-Aldrich, United States) were mixed on a rotating shaker overnight at 4°C. Unbroken cells were pelleted via centrifugation at 3,000 g for 5 min, and the supernatant was centrifuged at 40,000 g for 20 min. The pellet was then resuspended in 40 mL TEMB containing 4% IGEPAL CA-630 and centrifuged again at 40,000 x g for 20 min. The resulting pellet was then resuspended in 2 mL TEB + 10mM MgCl_2_ (TEMB) (5 mM Tris-HCL, pH 8.0, 1 mM EDTA, 10 mM MgCl_2_, 20 mM NaHCO_3_) and centrifuged at 5000 x g for 5 min before loading onto a 20-60% (v/v) sucrose gradient in TEMB buffer. Gradients were then centrifuged at 105,000 g for 60 min at 4°C; the milky band at the 40%-50% interface was collected, diluted in 10 mL TEMB buffer and centrifuged again at 105,000 g for 60 min. The final carboxysome pellet was then resuspended in 150 µL TEMB for the following structural and biochemical analysis.

Purified carboxysomes were first diluted to 8 mg mL^-1^. CaCl_2_, KCl, and MgCl_2_ stock solutions were prepared in TEMB buffer, with the Ca/K/Mg concentration ranging from 40 mM to 1000 mM, filtered and added to carboxysome samples at 1:1 ratio (v/v). The mixture was mixed gently and incubate at 30°C overnight. Protease Inhibitor Cocktail (Sigma-Aldrich, US) was added to reaction according to manufacture suggestions to avoid protein degradation.

### Negative staining electron microscopy

Negative staining electron microscopy was carried out as described previously ^39,42,43^. The carboxysomes (∼4 mg mL^-1^) were stained with 3% uranyl acetate on glow-discharged carbon-coated grids and then inspected with FEI 120 kV Tecnai G2 Spirit BioTWIN TEM equipped with a Gatan Rio 16 camera. Samples were visualized with ImageJ and statistically analyzed by Origin (OriginLab, US).

### SDS-PAGE analysis

SDS-PAGE analysis was performed following standard procedures. 10 μg purified carboxysomal proteins or 100 μg whole cell fractions were loaded per-well on 15% polyacrylamide gels and stained with Coomassie Brilliant Blue G-250 (ThermoFisher Scientific, UK).

### Cryo-EM SPA sample preparation and data collection

The *Cyanobium* sample was prepared by plunge-freezing in ethane onto the carbon side of Lacey ultra-thin carbon 400 mesh grids (Agar Scientific) using Vitrobot with a blotting time of 3.5s and blotting force of -15. The Grids were glow-discharged for 45s before use. Data were acquired with the Thermofisher 300kV Ttian Krios microscope equipped with a Falcon 4 direct electron detector with a Selectris energy filter operated with 10eV slit width. The pixel size is 1.171Å with a total electron dose of ∼40e^-^/Å^2^ for each frame movies. 13606 frame movies were acquired in total.

### Cryo-EM SPA data processing of Cyanobium carboxysome

For cryo-EM SPA of the *Cyanobium* α-carboxysomes, beam-induced motion was corrected using MotionCor2^44^ to generate dose-weighted micrographs from all movie frames. The contrast transfer function (CTF) was estimated using Gctf^45^. The particle picking, 2D and 3D classification and final refinement were conducted in Relion3.1^46^. The particles were automatically picked using a 2D class averages obtained from a subset of manual picked particles. The resulting particles were extracted at bin 4 and subject to several rounds of 2D classification and 3D classification with C1 symmetry, which resulted in a relatively clean dataset (6719 from 20982 particles). The final refinement with C1 symmetry resulted in a density map at a resolution at 38 Å, which is presented using ChimeraX^47^.

### Cryo-ET sample preparation and data collection

The purified *Halothiobacillus* α-carboxysomes were plunge-frozen in ethane onto lacey holy carbon grids (300 mesh, Agar Scientific) using Vitrobot or Leica GP2. The Grids was glow-discharged for 45s before plunge and gold fiducial beads (6nm) were mixed with the sample prior to sample application to grids. The excess solution was blotted with filter paper for 3 seconds with a humidity of 100% and temperature of 20 °C. The tilt-series were acquired using a ThermoFisher Titan Krios microscope operated at 300 keV, equipped with a K2 camera and Quantum energy filter in zero-loss mode with 20 eV slit width. The tilt-series were collected with SerialEM^48^ using dose-symmetric tilt-scheme starting from 0° with a 3° tilt increment by a group of 3 and an angular range of ±60°. The accumulated dose of each tilt series was around 120 e-/Å2 with a defocus range between -2 and -5 µm. Ten raw frames at each tilt were saved for each tilt-series. The Ca^2+^ treated *Halothiobacillus* α-carboxysomes and two *Cyanobium* α-carboxysomes datasets (apo and Ca^2+^ treatment) were collected with a K3 camera with SerialEM using similar parameters. Details of data collection are listed in Supplementary Data Table 1.

### Subtomogram averaging

Tilt-series from *Halothiobacillus* carboxysomes (apo and Ca^2+^ treatment) and Cyanobium carboxysomes (apo state) were aligned with IMOD using the gold fiducials, with the aid of in-house on-the-fly processing python script (https://github.com/ffyr2w/cet_toolbox). The center of each identified gold fiducial was manually checked. The Ca^2+^ treated Cyanobium carboxysome dataset do not have gold fiducials and were aligned using Aretomo^49^ in a fiducial-less way. Subtomogram averaging was performed using emClarity^32^. Rubisco crystal structure (PDB: 1SVD) was converted to density map at 20 Å resolution using *molmap* command in Chimera and subsequently used as the template for template matching in emClarity. Template matching was performed with 4x binned tomograms with a pixel size of 5.36 Å (hereafter bin4 tomograms) with or without ctf correction but filtered at the first zero of CTF (contrast transfer function) in emClarity. The resulting Rubisco coordinates were manually inspected to remove the false positives and the isolated Rubiscos outside carboxysomes. The Rubisco coordinates were also carefully checked against the bin4 tomograms to ensure that most of the Rubiscos inside carboxysomes are picked up. For *Halothiobacillus* carboxysomes (apo form), subtomograms from the first 60 tilt series (from 165 tilt-series) was used for subtomogram averaging and alignment. The averaging and alignment were firstly performed at bin3 with a pixel size of 4.02 Å for 4 cycles, bin2 (2.68 Å pixel size) for 8 cycles and bin1 for 4 cycles. We performed one round of tomoCPR at bin3 after bin1 alignment, and repeated the alignment at bin2 and bin1, which improved the overall density map. Duplicates of subtomograms were removed during alignment. The dataset was divided into two independent subsets during the alignment for a gold-standard metrics and the two subsets were combined in the final iteration, which resulted in the final resolution of 3.3 Å. C4 symmetry was applied throughout the alignment procedure, except the final 2 rounds of alignment using D4 symmetry. *Cyanobium* carboxysome dataset (apos state) were processed in the similar way without tomoCPR and the final density map is reconstructed using 2D tilt series images with cisTEM within emClarity package, at a resolution of 3.8 Å.

After the consensus alignment, Rubiscos from different positions from carboxysomes were extracted and reconstructed with cisTEM, with one round of local translational searches. Rubiscos from the three concentric layers in *Cyanobium* carboxysomes were selected based on radial distance distribution (Fig. 2c). Rubiscos within 300 Å distance from *Halothiobacillus* carboxysome center were extracted obtain a density map representing internal Rubiscos. The Rubisco along the *Halothiobacillus* carboxysomes shell were identified as following steps: the center of individual carboxysomes were manually labelled and further refined by averaging position of all the Rubiscos within the carboxysomes, and Rubisco within 400 Å were removed to only keep the Rubisco close to the shell. Since *Halothiobacillus* carboxysomes have various sizes and morphology, a further manual inspection of the remaining Rubisco coordinates was performed to remove the Rubiscos that are not along the shell.

### Identification of Rubisco string and subtomogram averaging

The Rubisco strings were obvious in the bin6 tomograms and can be identified from mapback coordinates after each of the Rubisco inside the carboxysomes was refined. Manual inspection was initially performed for a small dataset. We found Rubiscos in the string have their 4-fold axis along the string and most strings are organized in a similar orientation within the same carboxysome. For the large dataset, Rubisco in the string was identified by satisfying the following geometry restraints: (i) two tandem Rubisco in the string should have their 4-fold axis pointing the same or opposite direction, due to the D4 symmetry and (ii) the distance between the adjacent Rubiscos should be close to diameter of Rubisco. Manual inspection was performed to remove the Rubiscos that do not locate in the string. To obtain a map focusing on Rubisco interface, the center of Rubisco alignment box was shifted to the dimer interface along the string from Rubisco center and further few rounds of alignment were performed.

### Radial and angular distributions of Rubiscos

To calculate the radial and angular distribution of Rubiscos, the center of each carboxysomes were calculated as the average of all Rubiscos positions in each carboxysomes. The distance between each refined Rubisco and carboxysome center were calculated to generate radial distance distribution. A radial vector for each Rubisco was calculated pointing from the center of carboxysome to each Rubisco; the angle between the radial vector and 4-fold axis or Rubisco were calculated to generate radial anguar distribution.

### Model building and Refinement

Crystal structure (PDB 1SVD) of Rubisco was manually fit into the subtomogram averaging density map from *Halothiobacillus* carboxysome and further refined in Coot^50^ and Phenix.real_space_refine^51^. The structure of Rubisco subunits (CbbL and CbbS) from *Cyanobium* was initially predicted using AlphaFold2^33^ and rigid-body fit into the density map to generate the full structure (8CbbL and 8CbbS). The resulting structure were manually corrected in Coot before the molecular dynamics flexible fitting using Namdinator^52^. The surface electrostatic potential was calculated using APBS^53^ plugin in PyMOL. Calculations were performed at 0.15M ionic strength in monovalent salt, 298.15 K. Distribution and orientation of Rubiscos were presented in Chimera using *Place Object* plugin^56^ after converting emClarity metadata to the required format. The figures were prepared in Chimera^54^ and PyMOL^55^.

### Purification of H. neapolitanus Rubiscos and CsoS2-NTD-sfGFP

*H. neapolitanus* Rubisco expression vector *pAM2991-CbbLS-kanR* which constructed by inserting the coding sequence of *cbbL* and *cbbS* from *pHnCBS1D*^57^, into a pAM2991 vector that contain kanamycin resistance gene by Gibson assembly^58^ (NEBuilder® HiFi DNA Assembly). *E. coli* BL21(DE3) cells which contain pAM2991*-cbbLS-kanR* was grown at 37°C in LB broth that contain kanamycin at a final concentration of 50 μg mL^-1^ and induced with 1mM of IPTG when OD_600_ reaches 0.6 for overnight at 20°C. Cell lysates are obtained by sonication and CelLytic™ B Cell Lysis Reagent (Sigma-Aldrich, US). Cell debris are removed by centrifugation at 24,000 g and crude Rubiscos were obtained by ammonium sulfate precipitation. The crude Rubiscos were then loaded to a linear sucrose gradient (0.2M-0.8M) and centrifuged at 200,000 g for 4 hours. Sucrose layers were fractionated and then identified by SDS-PAGE and Rubisco containing fractions were load onto a HiTrap Q HP anion exchange chromatography column (Cytiva) by AKTA system. The eluent that contains pure Rubiscos were buffer-exchanged in dialysis tube to remove unwanted salt. Rubiscos were stored at 4°C for short-term or -80°C for long-term. *H. neapolitanus* CsoS2-NTD-sfGFP expression vector pCDF*-csoS2NTD-sfGFP* was designed as described previously^24^. The N-terminal 6xHis tagged N-terminal domain of CsoS2 were fused with super-folder GFP on C-terminus were inserted in the first cloning site of the pCDFDuet-1 vector (Novagen) by Gibson assembly^58^ (NEBuilder® HiFi DNA Assembly). Expression was performed as described above. Cell lysates were generated as described above, target proteins were purified with HisTrap HP (Cytiva) by AKTA system. Protein sequences are provided in Supplementary Table 2.

### In vitro Liquid–liquid phase separation assay and FRAP measurements

Purified *H. neapolitanus* Rubiscos and CsoS2-NTD-sfGFP were first diluted to 8 µM. A 100-μL reaction mixture was prepared by sequential addition of 80 μL sodium salt buffer (TEMB with 25 mM NaCl), 10 µL Rubisco, and 10 µL CsoS2-NTD-sfGFP. The mixture was mixed gently and applied to the center of uncoated Glass Bottom Dish 35 mm dish (ibidi) at volume of 20 μL. The mixture was then subject to incubation at 30 °C for 5 min. Salt treatment was accomplished by addition of equal volume calcium salt buffer (TEMB with 500 mM CaCl_2_) or sodium salt buffer as control. Droplets that rest on the bottom of the plate were captured by Zeiss LSM710/LSM780 Confocal Laser Scanning Microscope with 63x/1.30 oil objective. Fluorescent emission of GFP was captured with parameter described previously ^59^. FRAP experiments on formed condensates were performed as described previously with a 250 ms interval^60^.

## Supplementary Tables and Figures

**Supplementary Table 1.**
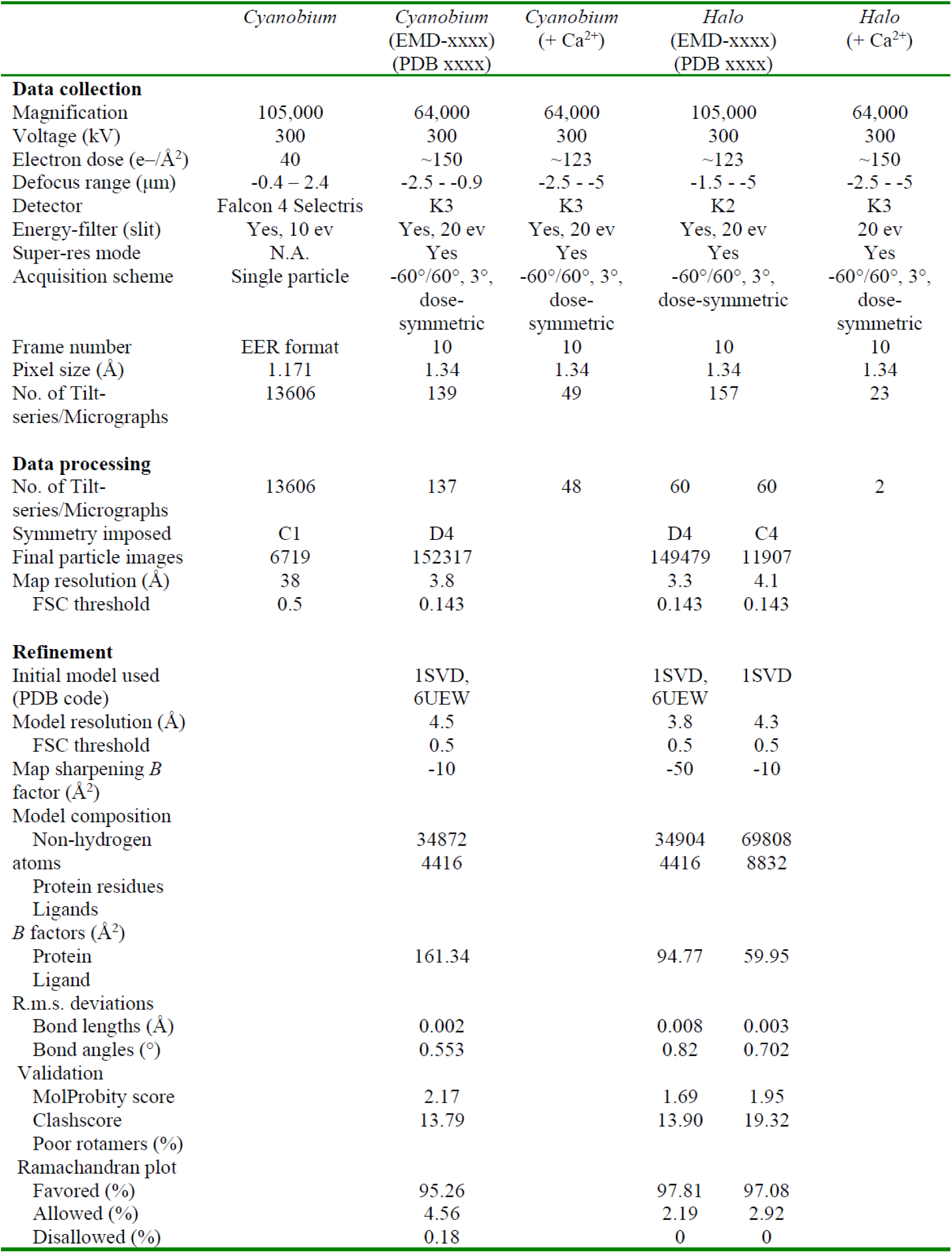
Data collection and model refinement statistics.

**Supplementary Table 2.**
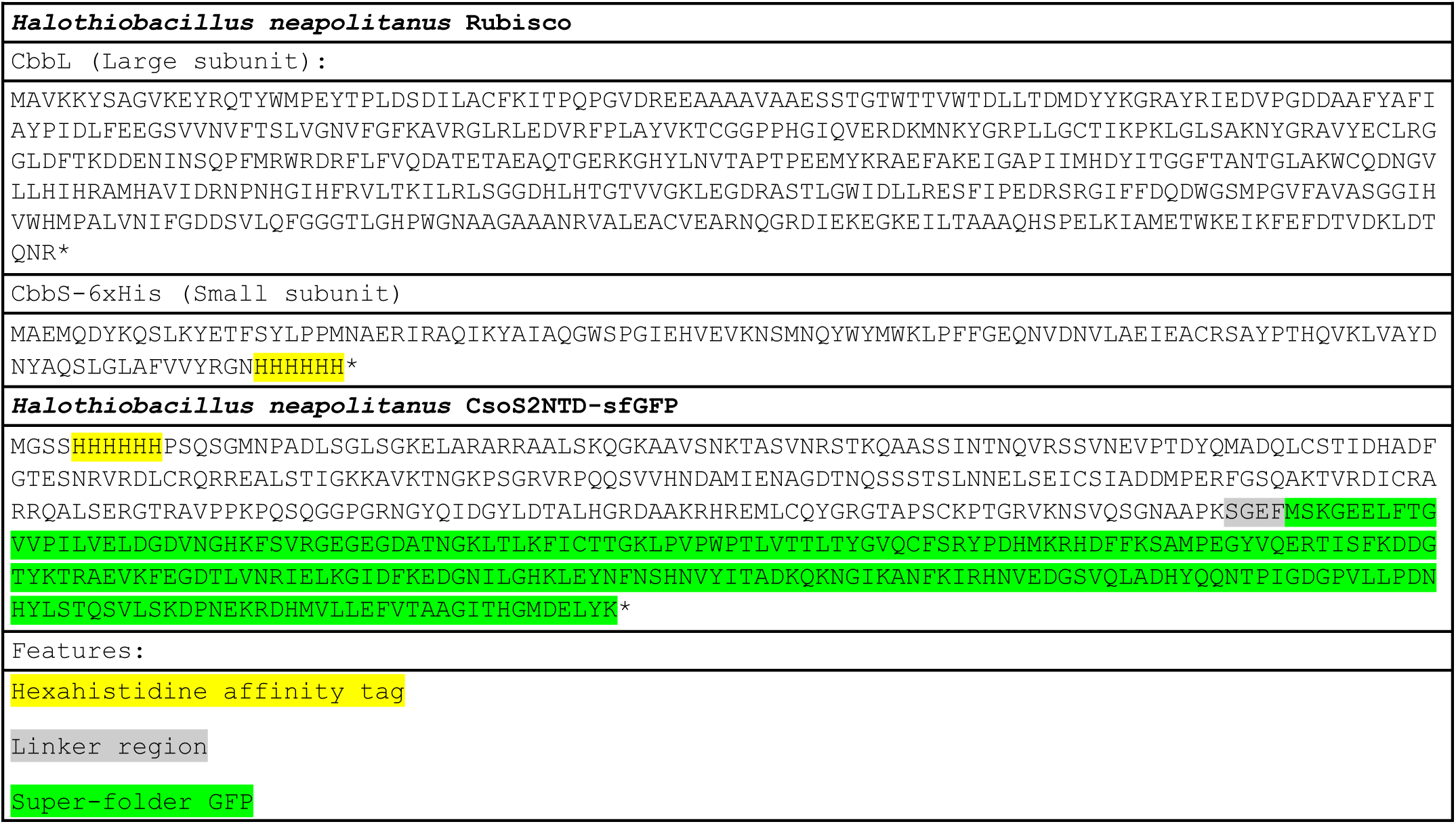
Protein sequences used in this study.

**Figure S1.**
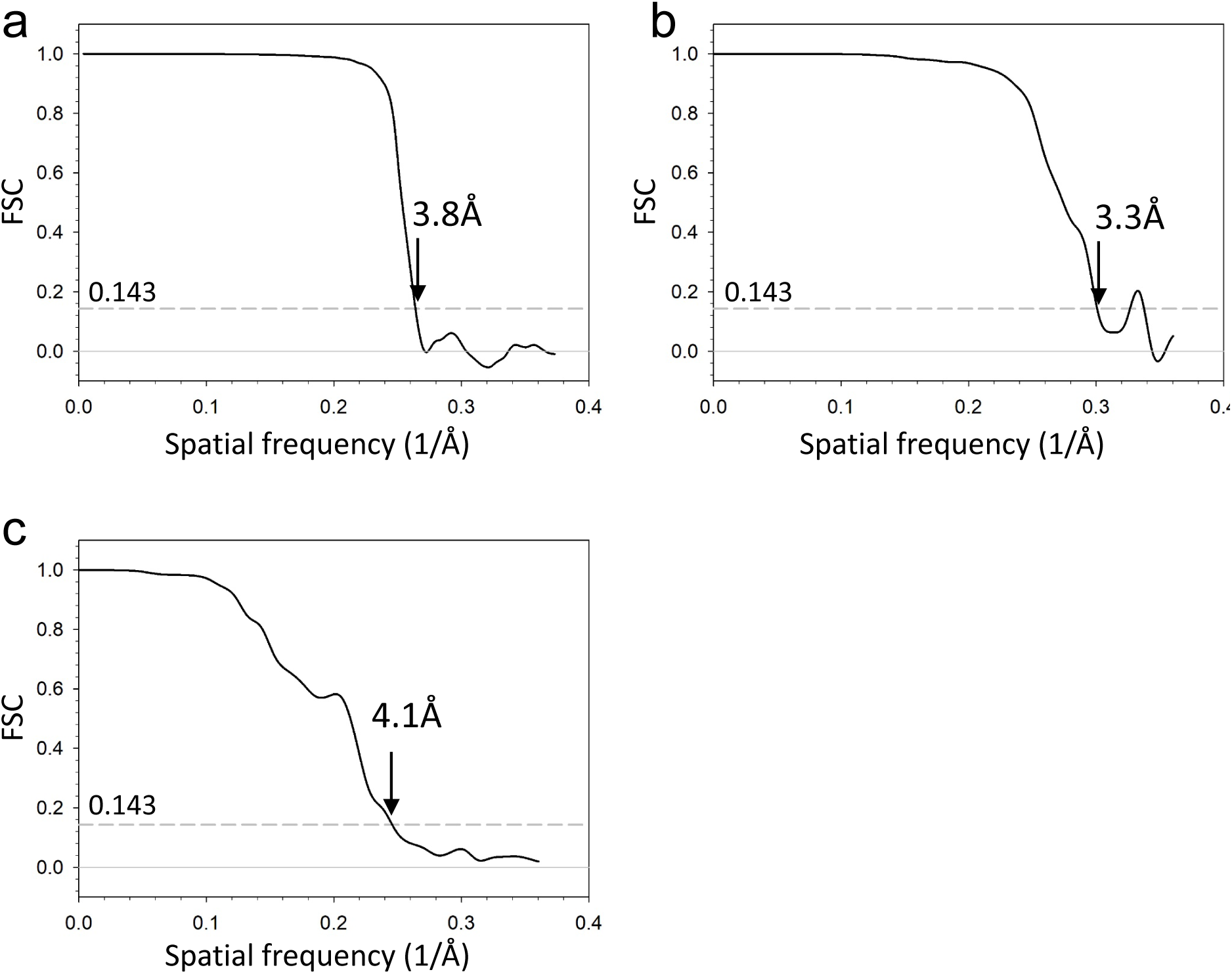
FSC plots of cryoET STA of *Cyanobium* and *Halo* α-carboxysomes. (a) FSC of Rubiscos STA map from *Cyanobium* carboxysomes. (b) FSC of Rubisco STA map from *Halo* carboxysomes. (b) FSC of Rubisco dimer STA map from *Halo* carboxysomes.

**Figure S2.**
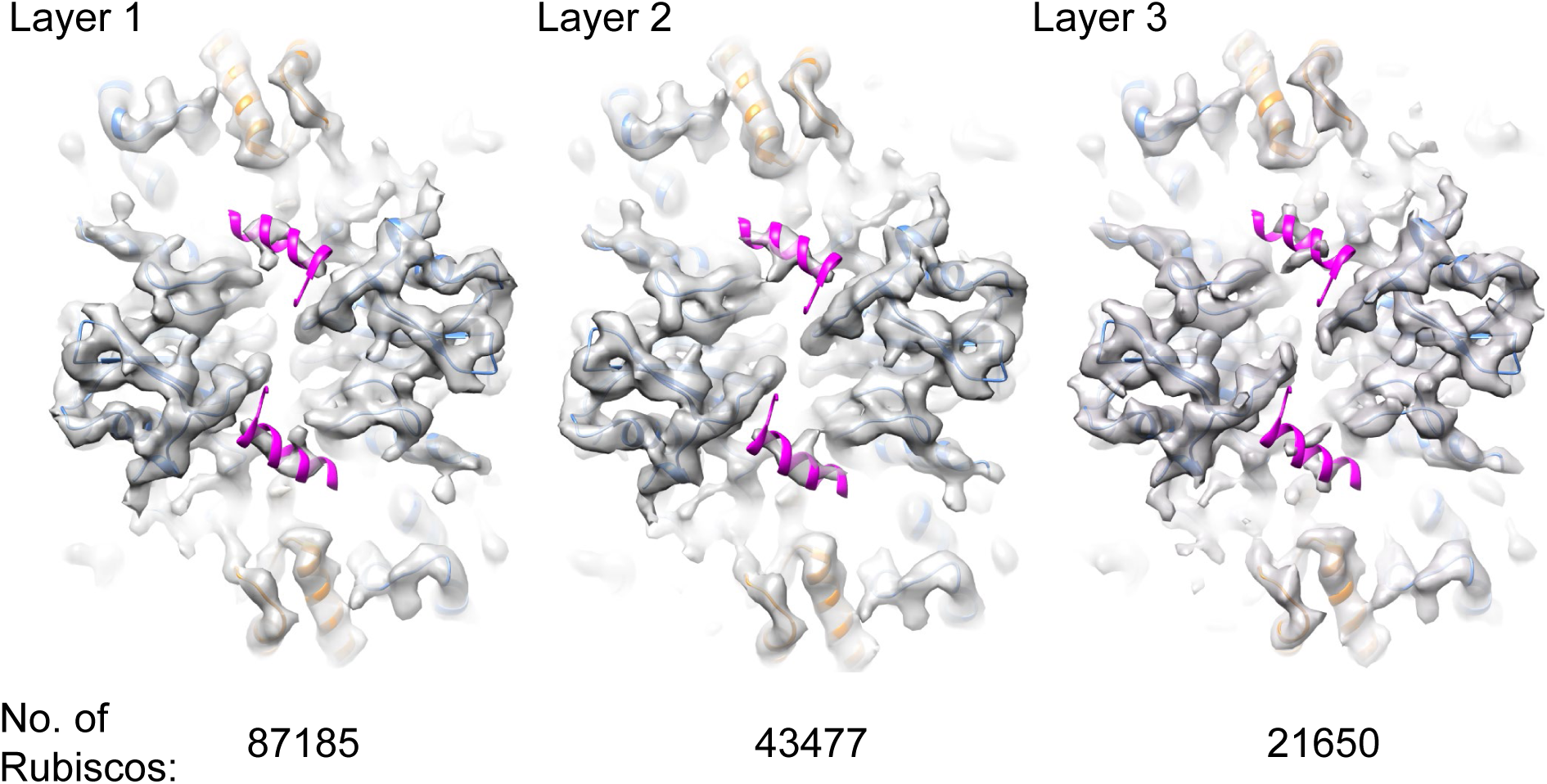
CsoS2 density in STA maps of Rubiscos in three concentric layer. Rubiscos from three layers were averaged, according to its radial distance distributions in Fig. 2c. From left to right: Layer 1: 350 - 600 Å, layer 2: 250 - 350 Å and layer 3: 0 - 250 Å. Numbers of subtomograms in each layer are listed below.

**Figure S3.**
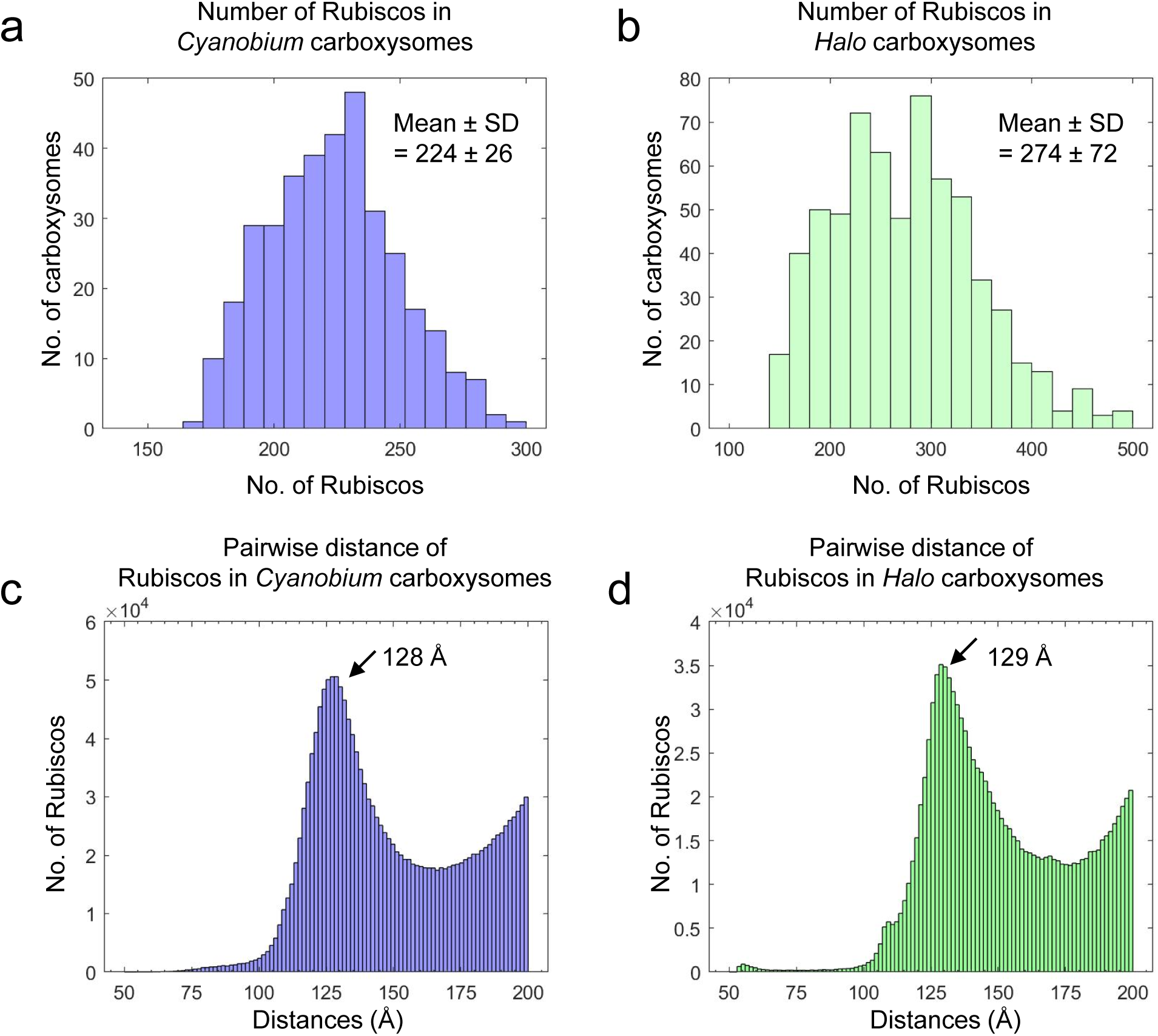
Comparison of *Cyanobium* and *Halo* carboxysomes. (a-b) Histogram of Rubisco numbers in two types of α-carboxysomes. Total numbers of carboxysomes are 360 in *Cyanobium* and 636 in *Halo*. Only the intact carboxysomes were included for Rubisco quantification. (c-d) Rubisco distances between pairs in two α-carboxysomes. Pairwise distances between two Rubiscos in each carboxysome were calculated and only distances within 200 Å were plotted. The distances between two neighbour RuBsiCos are peaked at 128 Å and 129 Å, respectively.

**Figure S4.**
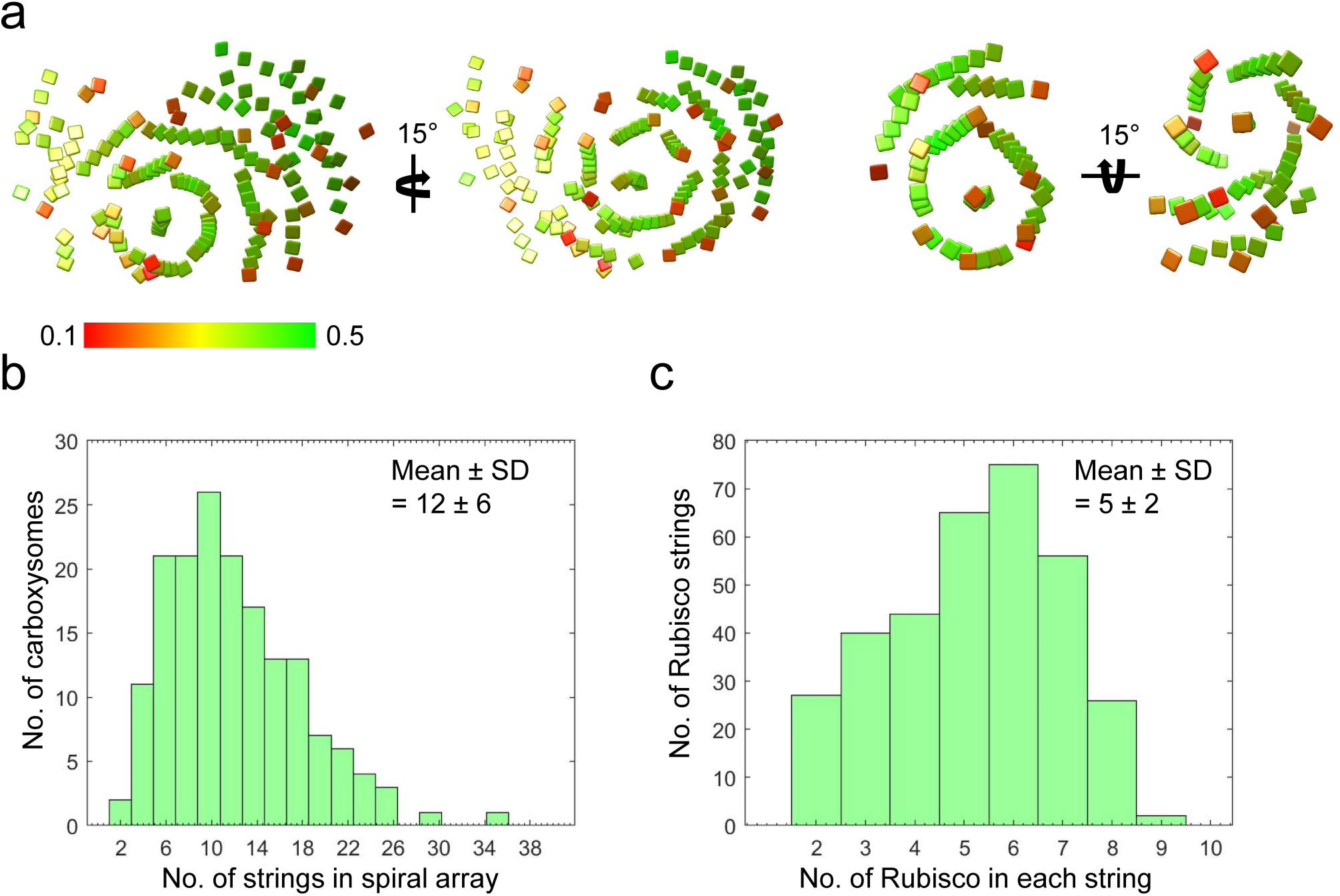
Characterization of Rubisco spiral arrays from *Halo* carboxysomes. (a) Two representative carboxysomes showing Rubisco spiral arrays in two orientation. The left carboxysomes contains 24 strings and the right one contains 10 strings. The position and orientation of individual Rubisco in the string is mapped back to the tomogram, shown as a discs normal the 4-fold symmetry axis of Rubisco and coloured according to the cross-correlation values between individual Rubisco and the STA map. (b) Histogram of Rubisco strings in *Halo* carboxysomes. The number of strings were quantified from the carboxysomes which show clear Rubisco arrays (167 out of 431 carboxysomes). (c) Histogram of Rubisco string lengths in *Halo* carboxysomes. Number of Rubiscos from 335 strings in 29 carboxysomes was counted.

**Figure S5.**
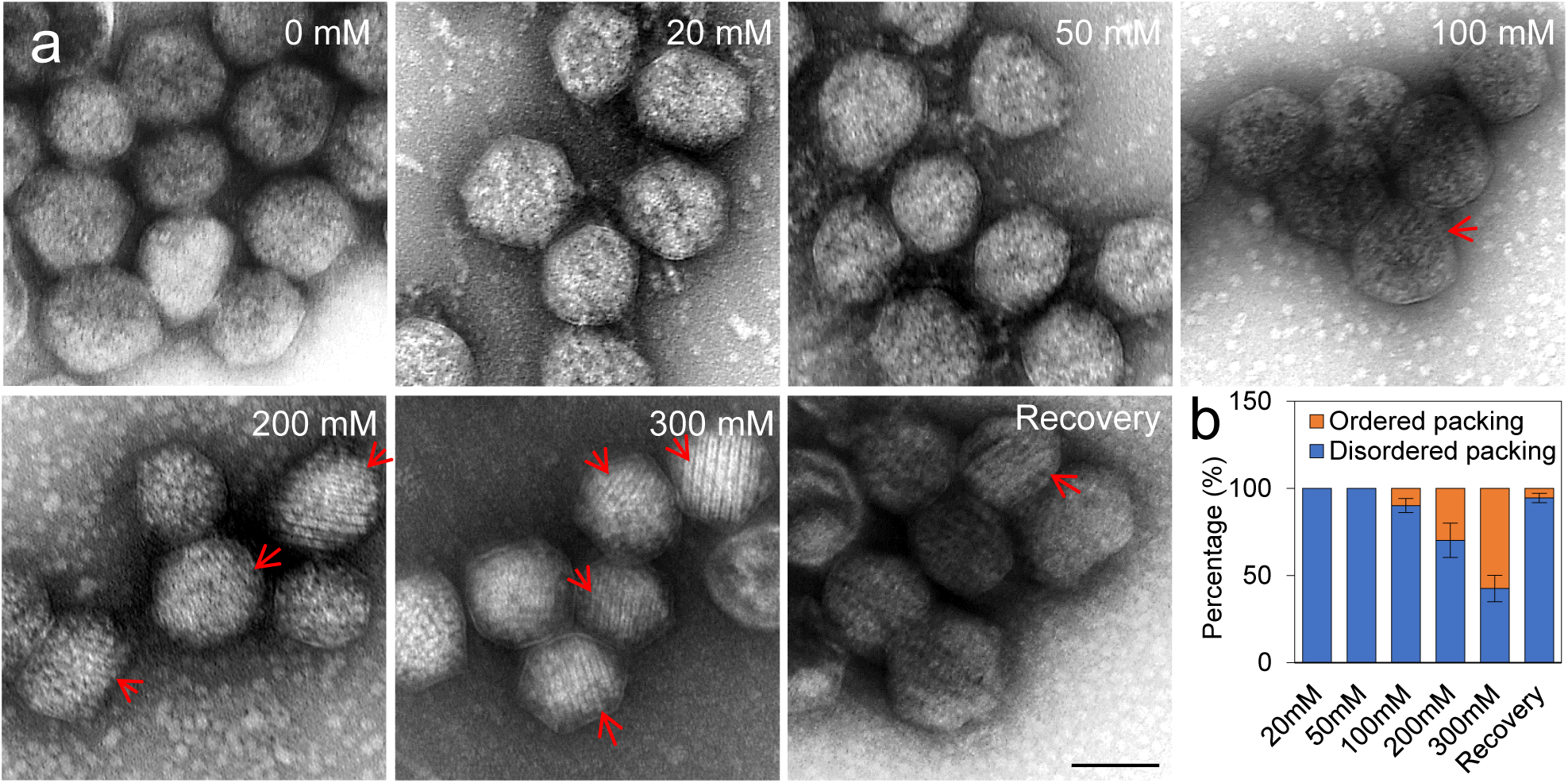
The effect of Ca^2+^ on *Halo* α-carboxysomes. (a) Negatively stained TEM images of *Halo* α-carboxysomes treated with Ca^2+^ at various concentrations and recovery from 300mM Ca^2+^ treatment. Red arrows point to string–containing carboxysomes. The concentrations of Ca^2+^ are indicated on top right of each treatment set. Scale bar =100 nm. (b) Quantification of α-carboxysomes containing Rubisco strings after Ca^2+^ treatment and recovery treatments (*n* = 100 as carboxysome counts for each treatment group averagely).

**Figure S6.**
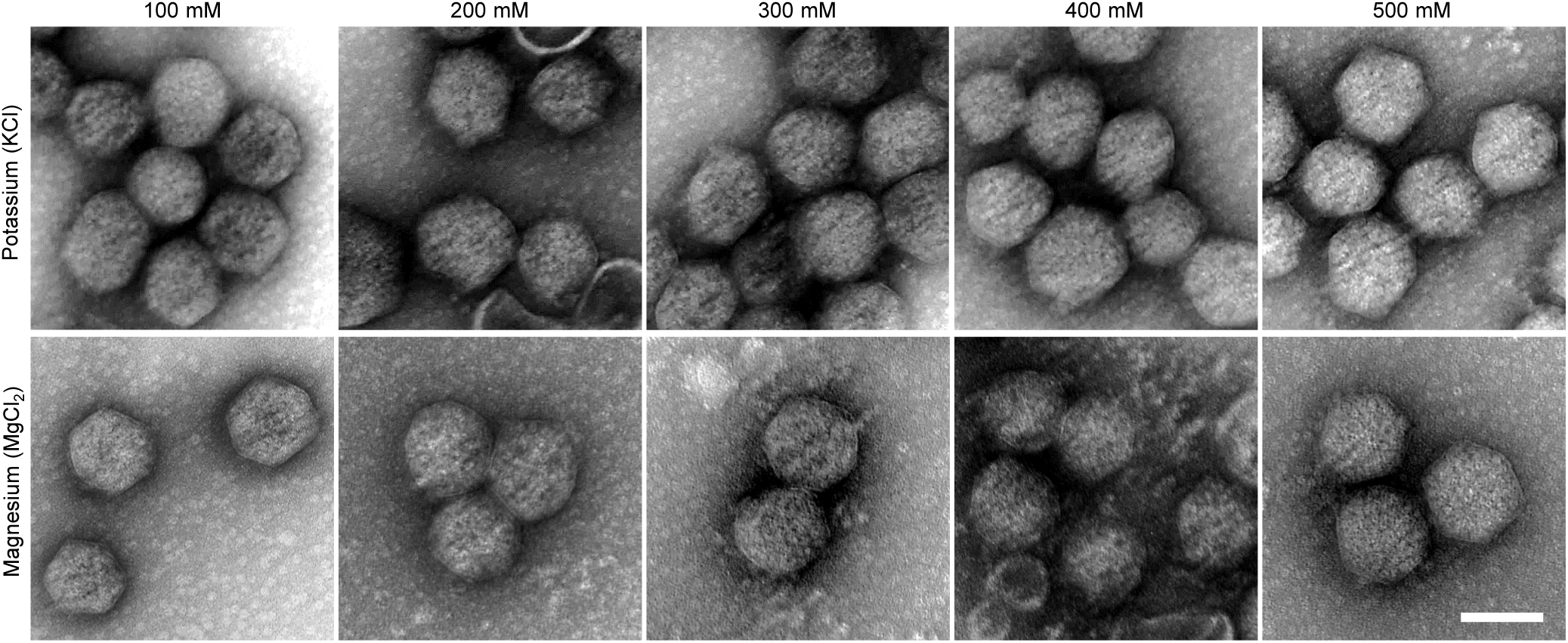
Potassium and magnesium treatment of *Halo* carboxysomes. Negatively stained TEM images of *Halo* α-carboxysomes treated with KCl (top) and MgCl_2_ (bottom) at the indicated salt concentrations. Scale bar = 100 nm.

**Figure S7.**
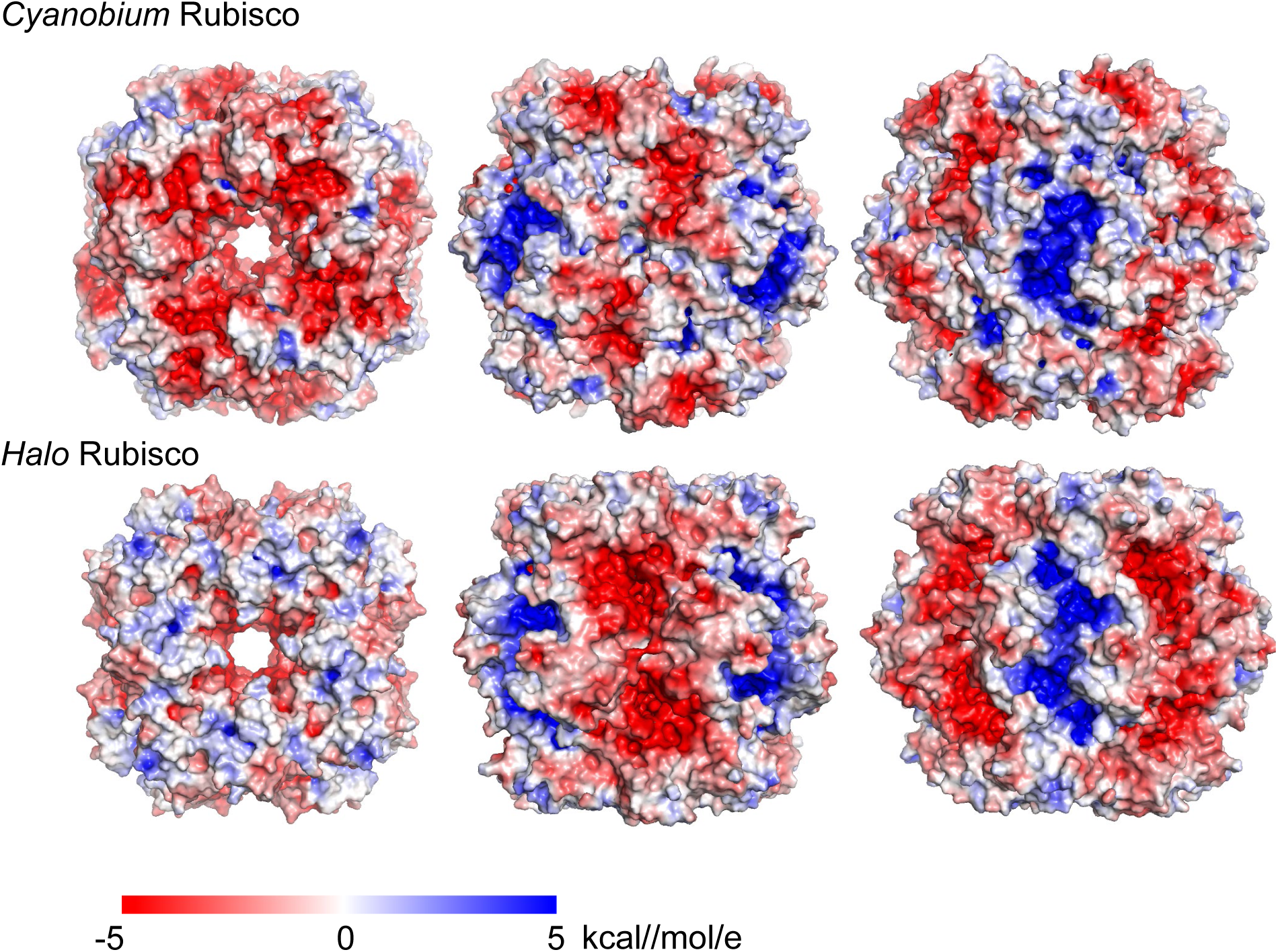
Surface electrostatic potential of Rubiscos from *Cyanobium* and *Halo*, presented in top view (left, along the 4-fold axis) and two side views (center and right, along the 2-fold axises). The surface electrostatic potential is calculated with APBS plugin in PyMOL. The potentials are on a [−5, 5] red-white-blue color map in units of kcal//mol/e.

## Notes

### Competing Interest Statement

The authors have declared no competing interest.

